# Gut-Associated Metabolites (GAMs) revitalize dysfunctional CD8⁺ T lymphocytes in Osteosarcoma

**DOI:** 10.64898/2026.07.30.741694

**Authors:** Leena Sapra, Asha Bhardwaj, Ankush Paladhi, Aasia Farhat, Taranjeet Kaur, Simran Preet Kaur, Chaman Saini, Venkatesan Sampath Kumar, Shah Alam Khan, Rupesh K. Srivastava

## Abstract

Osteosarcoma (OS) is one of the top ranking and deadliest primary malignant bone tumor of youngsters and adolescents. OS has poor prognostic features due to its immunosuppressive “*cold*” tumor microenvironment, obstructing the anti-tumor effector functions of immune cells (including cytotoxic CD8⁺ T lymphocytes)․ Recent developments have focused on the role of Gut Microbiota in modulating effector immune responses in various cancers. However, the immunomodulatory role of gut microbiota and its derived gut-associated metabolites (GAMs) in OS still remains unclear. Here we report that peripheral CD8⁺ T lymphocytes in osteosarcoma patients are less frequent in circulation, hypo-producers of effector cytokines (IFN-γ and TNF-α), and have compromised metabolic fitness․ We characterized and investigated the effect of a panel of microbiota-derived GAMs on CD8⁺ T lymphocytes, confirming their immunomodulatory role with respect to CD8⁺ T lymphocytes activation, cellular metabolism and effector functions․ Indole-3-lactic acid (ILA; product of tryptophan metabolism), was observed to be the strongest immunostimulant among all the GAMs studied. ILA enhanced the anti-tumor effector functions of CD8⁺ T lymphocytes through increased glucose uptake, a higher mitochondrial bio-mass and increased production of IFN-γ and TNF-α․ Moreover, ILA-primed cytotoxic T lymphocytes exhibited considerably higher cytotoxicity and increased apoptosis of osteosarcoma cells (U2OS)․ Altogether, our data demonstrates that ILA acts as a microbial-derived immune modulator to metabolically reprogram and restore dysfunctional CD8⁺ T lymphocytes against osteosarcoma․ This study for the first time demonstrates the therapeutic potential of exploiting the nexus between “*Gut-Immune-Bone Tumor”* ternary as a safe and cost-effective combinatorial immunotherapy against osteosarcoma․

**Graphical Abstract:** **Indole-3-Lactic Acid (ILA) Reinvigorates CD8^+^ T-Cell Immunity in Osteosarcoma**

Circulating CD8^+^ T cells from osteosarcoma patients exhibit impaired metabolic fitness, reduced effector cytokine production, and diminished tumoricidal activity. Screening of gut-associated metabolites identified the microbial tryptophan metabolite indole-3-lactic acid (ILA) as the most potent immunomodulator. ILA restores glucose uptake and mitochondrial biomass, enhances IFN-γ and TNF-α production, promotes polyfunctional CD8^+^ T-cell responses, and significantly improves CTL-mediated killing of osteosarcoma cells, highlighting its potential as a microbiota-derived immunometabolic therapeutic for osteosarcoma.

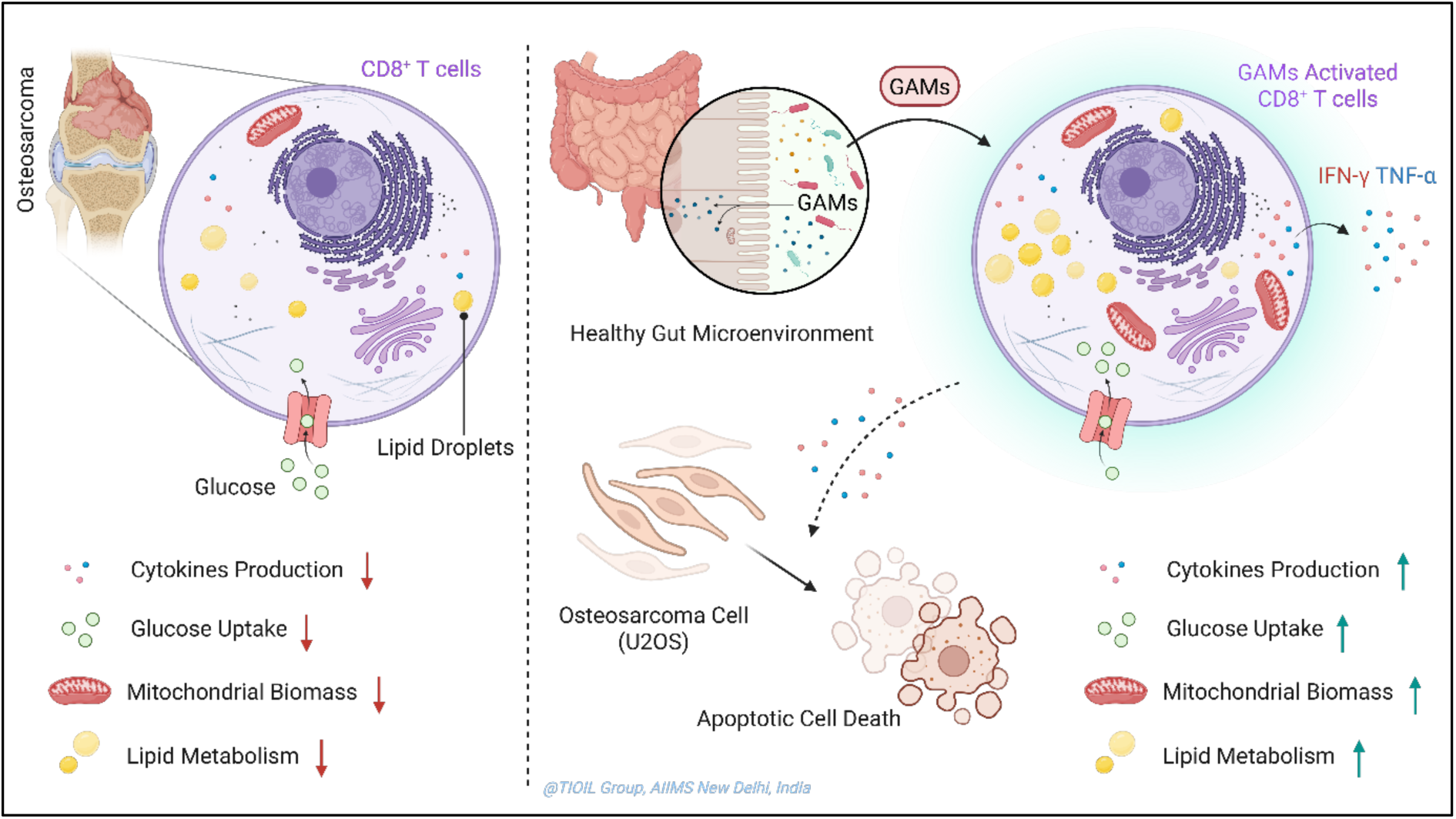

## 1.0 Introduction

Osteosarcoma (OS) is one of the top-ranking deadliest skeletal malignancy of youngsters and adolescents, accounting for around 20% of all primary skeletal malignancies and showing a bimodal age distribution with a major risk of incidence peak during adolescence and a minor one in the elderly^1,2^. In India, it is one of the leading primary paediatric malignancies (∼11% of the total) with an abruptly increasing rate of incidence per year^3^. Osteosarcoma is an osteoid-forming malignancy of mesenchymal origin that primarily developed in the metaphysis of long bones^1^. In spite of, notable progresses in multidimensional disease-management strategies, including surgical resection, neoadjuvant-based chemotherapy, advanced immunotherapy, the prognostic betterment in terms of disease-free overall survival of patients with recurrent or metastatic disease has remained largely unimproved since several past decades^4^. The unsatisfactory clinical outcome due to prompt pulmonary metastasis, intrinsic or acquired chemoresistance, tumor recurrence and treatment-associated co-lateral toxicities, spotlighting the urgent need to explore a novel, more effective and less toxic treatment strategies^4,5^.

Overall, osteosarcoma can be considered as an immunosuppressive tumor due to its notably high “cold” microenvironment, which promotes dysfunction of cellular and molecular effector anti-tumor immunity^6^. Among the all-immune cell populations, mainly CD8^+^ cytotoxic T lymphocytes (CTLs) are considered as the principal tumoricidal mediators via secreting tumor cell eliminating inflammatory and/or cytotoxic effector cytokines, including tumor necrosis factor-α (TNF-α), interferon-γ (IFN-γ), perforin-1 and granzyme-B^7^. However, immunoregulatory checkpoint pathway activation, persistent exposure of tumor associated antigens (TAAs), and metabolic challenges within the hypoxic and nutrients limited tumor microenvironment trigger CD8^+^ T cell dysfunction, resulting in dampened anti-tumor effector cytokines secretion, diminished cytotoxicity and rapid exhaustion of tumor infiltrating T lymphocytes^8,9^. Emerging investigations by multiple independent research group further revel that T lymphocytes activation state and effector functionality are critically coupled with cellular metabolic fitness, including adequate carbohydrate (mainly glucose) utilization, mitochondrial biogenesis and integrity and lipid metabolism^10,11^. Consequently, metabolic deficiency has emerged as a primary contributor to insufficient anti-tumor immune responses of CD8^+^ T cell in osteosarcoma^12^.

Alongside other research groups our group has also spent more than a decade to decode the *“Gut-Immune-Bone”* nexus. We have extensively investigated the immunomodulatory potential of gut microbiota and also microbiota-derived GAMs (including SCFAs, secondary bile acids and tryptophan metabolites) in maintaining homeostatic skeletal health ^13–19^. These gut-microbiota-derived metabolites regulate the activation, cellular differentiation and bioenergetics of immune cells of different lineages, thereby reprogramming both innate and adaptive arms of systemic immunity^20^. Emerging evidences claimed that specific GAMs augment the effector immune function of cytotoxic CD8^+^ T lymphocytes by fine-tuning cellular metabolism. Consequently, upregulate IFN-γ and TNF-α secretion, enhancing cytotoxic potential, improving anti-tumor immune responses, and widening immunotherapeutic spectrum^21,22^. Moreover, promising immunomodulatory properties of gut microbiota-derived indole metabolites, especially indole-3-lactic acid (ILA), has been emerged to upregulate T cell-mediated anti-tumor immunity through delicate metabolic and epigenetic remodelling^23,24^. Collectively, these findings emphasize the potential of gut-associated metabolites as emerging immunomodulators for improving T-cell responses against growing tumors.

Our group has established the *“Gut-Immune-Bone”* nexus by decoding the immunomodulatory role of gut-microbiota and their metabolites in maintaining homeostatic bone health^15–18^. More recently, this idea has been extended to bone tumor by portraying the interplay between the gut microbiome, intratumoral microbiota, and immunosuppressive tumor microenvironment in skeletal malignancy, providing a prominent conceptual framework for exploiting the role of gut-associated metabolites as a novel immune therapeutic modulator in bone cancers^19^. Building upon these foundations, we hypothesized that gut-associated metabolites may enhance effector immune functions and anti-tumor immunity of CD8^+^ cytotoxic T lymphocytes in osteosarcoma.

Regardless of significant growth in the research field of gut-associated metabolites in regulating immune responses to cancer, their role in modulating CD8^+^ cytotoxic T lymphocytes, especially reverting dysfunctional state and reprogramming cellular energetics in osteosarcoma remains mostly unrevealed^21,25^. We therefore hypothesized that selected gut-associated metabolites may reprogram the anti-tumoral functionality as well as metabolic fitness of CD8^+^ T lymphocytes in osteosarcoma. To address this, we first characterized the overall frequency, effector function and metabolic status of circulating CD8^+^ T lymphocytes within the total PBMCs from osteosarcoma patients. We further assessed a wide range panel of gut-associated metabolites to identify most competent candidate for rejuvenating anti-tumor effector functionality and immunometabolic fitness of dysfunctional CD8^+^ T lymphocytes.

Finally, we identified the effects of the most potent metabolite, indole-3-lactic acid (ILA), on T lymphocytes-mediated cytotoxicity against osteosarcoma cells (U2OS). By integrating patient-derived immune profiling with functional and metabolic analyses, this investigation pinpoints ILA as an emerging immunomodulator for triggering CD8^+^ T-cell-mediated tumoricidal responses, providing a prominent rationale for the future development of gut metabolite-based novel combinatorial chemo-immunotherapeutic strategies for prognostic betterment in osteosarcoma.

## 2.0 Materials and Methods

### 2.1 Study design

The study, focused on investigating the functional and metabolic status of the circulating CD8^+^ T lymphocytes in osteosarcoma (OS) patients and the immunomodulatory activity of gut-associated metabolites (GAMs). Flow cytometry, was employed to analyze the frequency, effector cytokine secretion and metabolic fitness of CD8^+^ T lymphocytes in both OS patients and healthy control. Next representative GAMs were screened for their ability to modulate the anti-tumor immune response of CD8^+^ T lymphocytes. Subsequently, the effect of indole-3-lactic acid (ILA; the most potent metabolite), was studied on CD8^+^ T lymphocytes metabolism and CTL-mediated lysis of osteosarcoma cells under *in-vitro* co-culture conditions.

### 2.2 Human subjects and ethical approval

Blood samples were collected after informed consent from ten osteosarcoma patients (histologically confirmed) and ten healthy volunteers of similar age group. Only treatment-naïve OS patients, between the age of 8 to 25 years, irrespective of their metastatic status were included in the current study. Moreover, OS patients with other malignancies or already experienced chemotherapy, pre-existing autoimmune disease, history of chronic inflammatory disorders or any major systemic comorbidities were excluded. Blood samples were harvested in heparinized vacutainers and processed immediately for PBMC isolation for further experimentation. The present study was approved by the Institutional Ethics Committee of the All-India Institute of Medical Sciences (AIIMS), New Delhi (Approval No. AIIMSA2271) as per Declaration of Helsinki.

### 2.3 Reagents, antibodies and gut-associated metabolites

Fluorochrome-conjugated primary antibodies against CD3 and CD8 T cell surface markers; TNF-α and IFN-γ effector cytokines; and Annexin V for detecting cellular apoptosis was procured from eBiosciences (USA). Metabolic probes, 2-NBDG, MitoTracker Deep Red FM, and BODIPY 493/503, PMA, ionomycin, monensin, HIAA, ILA, DCA, 3-Oxo-LCA, 7-Keto-LCA, AILCA, acetate, butyrate, propionate, valerate, RPMI-1640, DMEM high glucose, FBS was procured from Thermo Fisher Scientific (USA). Lymphoprep was procured from Serumwerk (Germany), and human CD8^+^ T cell negative selection kit was procured from BD Biosciences (USA). All the reagents and kits were used in accordance with the recommended protocols from the corresponding companies.

### 2.4 Isolation of peripheral blood mononuclear cells (PBMCs)

Using Lymphoprep, PBMCs were isolated from heparinized blood by density-gradient centrifugation. Blood was mixed with an equal volume of sterile PBS, layered over Lymphoprep and centrifuged under brake less conditions at 800 x g for 25 minutes. The PBMC layer was removed and washed 2x, resuspended in 10% FBS containing RPMI-1640 complete cell culture medium. The number and viability of the isolated PBMCs were assessed by trypan blue exclusion assay prior to further analysis.

### 2.5 Isolation of CD8⁺ T lymphocytes

CD8^+^ T lymphocytes were separated from freshly isolated PBMCs via magnetic selection in accordance with the manufacturer’s protocol to separate untouched CD8^+^ T lymphocytes from total PBMCs. Briefly, PBMCs were incubated with a cocktail of biotinylated antibodies, followed by streptavidin-conjugated magnetic particles, and the unlabelled cell fraction was collected, washed, resuspended in 10% FBS containing complete RPMI-1640 cell culture medium. The purity of isolated CD8^+^ T cells was routinely found to be > 95% as confirmed by flow cytometry.

### 2.6 PBMC stimulation and intracellular cytokine staining

Isolated PBMCs (1x10^6^ cells per well) were cultured in 10% FBS containing RPMI-1640 complete cell culture medium, at 37°C in a humidified incubator maintaining 5% CO_2_ constantly. Further, for *in-vitro* stimulation of cytokine production, cultured PBMCs were pulsed with PIM [a mixture of 50 ng/mL of PMA (phorbol 12-myristate 13-acetate), 1 µg/mL of ionomycin and 1 μg/mL of monensin] for 5 hours. Monensin was used to inhibit extracellular release of cytokines during the course of pulsation period. Following pulsation, cells were washed with sterile PBS to remove PIM, fixed in 4% PFA containing ice-cold PBS for 10 minutes and permeabilized (Transcription Factor Staining Buffer Set), and finally stained with distinct fluorochrome-conjugated anti-human primary antibodies against distinct surface as well as intracellular markers (CD3, CD8, IFN-γ, and TNF-α). Stained samples were finally acquired on a BD FACSymphony A5 flow cytometer and analyzed using FlowJo software (v10.8).

### 2.7 Treatment with gut-associated metabolites (GAMs)

PBMCs were cultured by following the same way mentioned earlier. Thereafter, treated with equal non-cytotoxic concentration (10 µM) of ten different representative GAMs, including HIAA, ILA, DCA, 3-Oxo-LCA, 7-Keto-LCA, AILCA, acetate, butyrate, propionate, and valerate for 24 hours. Cisplatin (8 µM) was included as the reference treatment. Following 24 hours incubation, cells were treated with monensin (5 hours prior to termination) to trap intracellular cytokines to facilitate detection via flow cytometry.

### 2.8 Assessment of metabolic fitness

The metabolic status of CD8^+^ T cells were evaluated by measuring intracellular glucose uptake, overall mitochondrial bio-mass and intracellular lipid content using fluorescent probe. 150 µM working concentration of 2-NBDG, 100 nM of MitoTracke Deep Red FM and 1 µg/mL of BODIPY 493/503 was used to detect glucose uptake, mitochondrial bio-mass and lipid content respectively. Following treatment, PBMCs were incubated with the metabolic probes along with anti-human CD8 antibody for 30-45 minutes at RT, washed with chilled PBS and acquired on a BD FACSymphony A5 flow cytometer. Flow cytometric data was analyzed using FlowJo software (v10.8). Confocal microscopy was further done to visualize overall mitochondrial bio-mass and intracellular lipid droplets in selected groups.

### 2.9 Cell culture

The human osteosarcoma cell line U2OS (ATCC HTB-96) was maintained in 10% FBS containing DMEM complete cell culture medium, at 37°C in a humidified incubator maintaining 5% CO_2_. Cells were routinely passaged at around 60-70% confluency by detaching from the surface of tissue culture flask using 0.25% trypsin-EDTA solution, followed by washing with sterile PBS and were used for experiment when the new inoculums reached logarithmic growth phase.

### 2.10 CD8⁺ T lymphocytes co-culture with U2OS

For functional and co-culture cytotoxicity assays, freshly isolated CD8^+^ T cells were cultured in complete RPMI medium in the presence or absence of indole-3-lactic acid (ILA; 10 µM) for 24 hours at 37°C in a humidified incubator maintained at 5% CO_2_. For intracellular cytokine analysis, cells were stimulated with monensin, 5 hours prior to staining. For cytotoxicity assays, ILA primed CD8^+^ T cells (without monensin) were used in co-culture experiments with U2OS osteosarcoma cells at different ratios.

### 2.11 CTL-mediated cytotoxicity assay

To estimate the cytotoxic (tumoricidal) potential, ILA-primed CD8^+^ T cells were co-cultured with U2OS osteosarcoma cells (seeded in 24-well plates with around 50% confluency) at different effector-to-target (E:T or CD8^+^ T cells: U2OS) ratios (1:1, 5:1, and 10:1) for 24 hours under standard culture conditions mentioned earlier. Following incubation, apoptosis of U2OS cell population was measured by Annexin V/PI counter staining, followed by flow cytometric analysis. During flow cytometric acquisition and analysis, CD8 positive populations were excluded by gating for measuring apoptosis in only target cells (U2OS) within heterotypic co-culture samples. Data were analyzed using FlowJo software (v10.8).

### 2.12 Flow cytometry

All Flow cytometry based studies were executed by using a BD FACSymphony™ A5 flow cytometer (BD Biosciences). Cell-surface marker staining was performed using distinct fluorochrome-conjugated anti-human primary antibodies against different markers, mentioned earlier. Whereas, cytosolic cytokine staining was carried out by fixation and permeabilization followed by staining with distinct fluorochrome-conjugated anti-human primary antibodies against different specific cytokines. For apoptotic assays, Annexin V-FITC and propidium iodide (PI) counter staining was performed in addition with fluorochrome-conjugated anti-human CD8 antibody-mediated surface staining to differentiate apoptotic target cells (U2OS) from effector cells (ILA-primed/unprimed CD8^+^ T cells). Data were analyzed using FlowJo software (v10.8).

### 2.13 Confocal microscopy

Using MitoTracker Deep Red FM and BODIPY 493/503, mitochondrial bio-mass and intracellular lipid droplet were visualized via confocal laser scanning microscopy. For each experimental group, fluorescence images were obtained under identical imaging conditions. The images were analyzed via the ImageJ (Fiji) image analysis software.

### 2.14 qRT-PCR

Gene expression studies were executed using quantitative real-time PCR (qRT-PCR) on a CFX Opus 96 Real-Time PCR System (Bio-Rad, USA). Initially, total cellular RNA was extracted from ILA/cisplatin-primed CD8^+^ T lymphocytes from OS patients. Therefore, reverse-transcribed to form cDNA, followed by amplified using selected effector cytokines (mainly IFN-γ, TNF-α, Perforin-1 and Granzyme-B) gene-specific primers with 2x SYBR Green Master Mix (Promega, USA). GAPDH gene expression status was used as the endogenous reference. Relative gene expression was executed as a value of delta threshold cycles (ΔCT).

### 2.14 Statistical analysis

All experiments were performed using at least three or more independent biological replicates. Results are represented as mean values with the corresponding SD of the mean (±SEM). Statistical analyses were executed with the help of GraphPad Prism (Version 10.0) software. Statistical comparisons between experimental groups were carried out using paired or unpaired Student’s t-test or one-way ANOVA, as per the compatibility to experimental design. In all cases the p value ≤ 0.05 was considered as statistically significant. Significance denoted as *p ≤ 0.05, **p ≤ 0.01, ***p ≤ 0.001, and ****p ≤ 0.0001.

## 3.0 Results

### 3.1 CD8⁺ T cells from OS patients display impaired effector function

To assess the functional state of circulating CD8^+^ T cells in osteosarcoma (OS) patients, PBMCs (Peripheral Blood Mononuclear Cells) isolated from OS patients and healthy control were pulsed with PIM (PMA-Ionomycin-Monensin) and analysed by flow cytometry for CD8^+^ T-cell prevalence and intracellular yield of effector cytokines (IFN-γ and TNF-α) (**Figure 1a**). Interestingly, OS patients displayed a significant decline in the prevalence of circulating CD8^+^ T cells compared with healthy control (**Supplemental Figure S1a**). More importantly, CD8^+^ T cells from OS patients yielded significantly lesser levels of IFN-γ and TNF-α following PIM stimulation (**Figure 1b–f**), pointing a marked impairment in their effector function. Given the crucial contributions of these cytokines in cytotoxic T-cell-mediated anti-tumor immune responses, their dampened production infers systemic immune dysfunction, thereby enhancing immune evasion and disease progression in OS. Altogether, these findings establish that circulating CD8^+^ T lymphocytes from OS patients have primarily impaired effector function, insisting further investigation into the mechanistic detailing behind this dysfunction.

**Figure 1.**
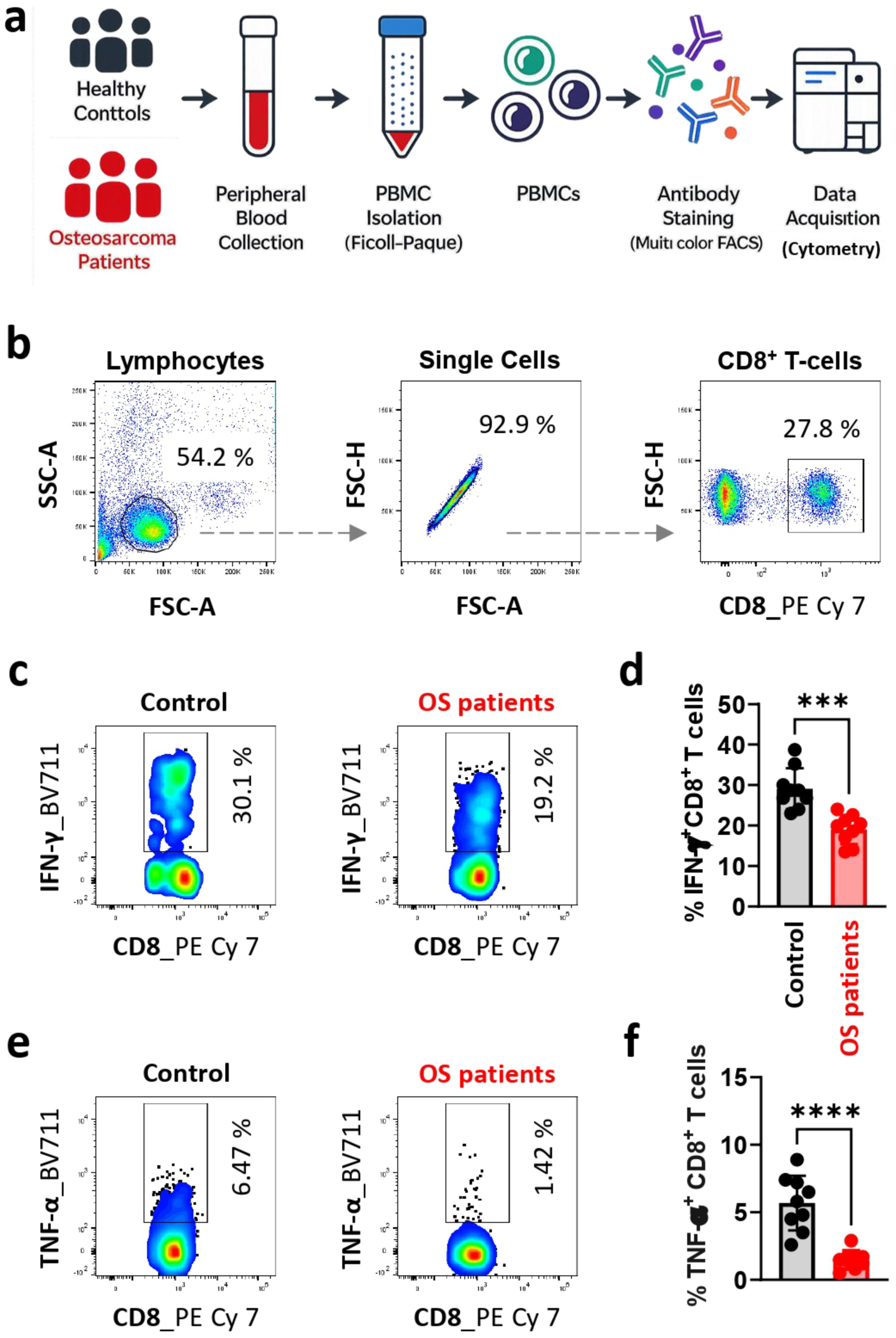
Circulating CD8^+^ T cells from osteosarcoma (OS) patients display impaired effector function. **(a)** Experimental workflow. Peripheral blood mononuclear cells (PBMCs) were isolated from healthy controls and OS patients, stimulated with PMA (50 ng/mL) and ionomycin (1000 ng/mL) for 5 h, followed by intracellular staining for IFN-γ and TNF-α and flow cytometric analysis. **(b)** Gating strategy for identification of CD8^+^ T cells. **(c, e)** Representative pseudocolor contour plots and **(d, f)** quantification of IFN-γ^+^ and TNF-α^+^ CD8^+^ T cells, respectively, in healthy controls and OS patients. Data are presented as mean ± SEM (n = 6 per group). Statistical significance was determined using an unpaired Student’s t-test. p ≤ 0.05 was considered significant; *p ≤ 0.05, **p ≤ 0.01, ***p ≤ 0.001, and ****p ≤ 0.0001.

### 3.2 CD8⁺ T cells from OS patients display compromised metabolic fitness

Effector functions of T cells are closely coupled to cellular metabolism; thus, we next looked into the metabolic status of circulating CD8^+^ T cells from OS patients. Glucose uptake, mitochondrial bio-mass, and intracellular lipid content were examined using the fluorescent probes 2-NBDG, MitoTracker Deep Red FM, and BODIPY 493/503, respectively (**Figure 2a**). In comparison with healthy controls, isolated CD8^+^ T lymphocytes from OS patients displayed considerably downregulated glucose uptake, hinting impaired cellular glycolytic pathway (**Figure 2b,c**). In addition, mitochondrial bio-mass was noticeably diminished, as confirmed by both flow cytometric and confocal imaging analyses (**Figure 2d,e,g**). Intracellular neutral lipid content was likewise significantly dampened (**Figure 2f,g**), intimating compromised metabolic arsenals. Collectively, these observations uncover the state of metabolic dysfunction in circulating CD8^+^ T cells from OS patients, which may volunteer to their compromised effector immune responses against osteosarcoma.

**Figure 2.**
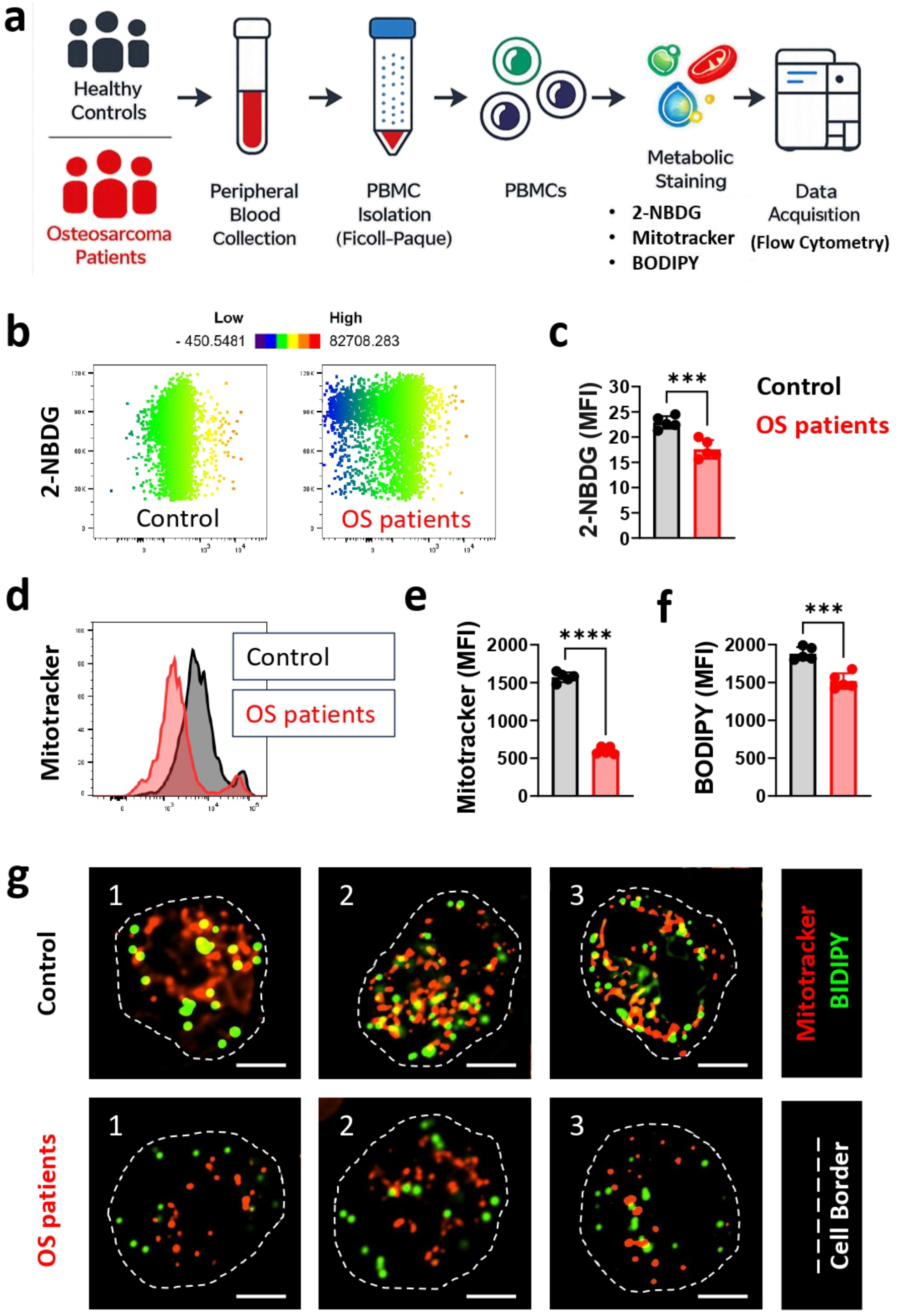
Circulating CD8^+^ T cells from osteosarcoma (OS) patients exhibit impaired cellular metabolism. **(a)** Experimental workflow. Peripheral blood mononuclear cells (PBMCs) from healthy controls and OS patients were stimulated with PMA (50 ng/mL) and ionomycin (1000 ng/mL) for 5 h, followed by staining with 2-NBDG, MitoTracker, and BODIPY for flow cytometric analysis. **(b, c)** Representative flow cytometry plots (heatmap dot plot) and quantification of glucose uptake (2-NBDG) in CD8^+^ T cells. **(d, e)** Representative histograms and quantification of mitochondrial biomass (MitoTracker). **(f)** Quantification of lipid accumulation (BODIPY) in CD8^+^ T cells. **(g)** Representative fluorescence microscopy images (three independent fields per group) showing mitochondrial biomass (MitoTracker, red) and lipid droplets (BODIPY, green) in CD8^+^ T cells from healthy controls and OS patients. Data are presented as mean ± SEM (n = 6 per group). Statistical significance was determined using an unpaired Student’s t-test. p ≤ 0.05 was considered significant; *p ≤ 0.05, **p ≤ 0.01, ***p ≤ 0.001, and ****p ≤ 0.0001.

### 3.3 GAMs restore IFN-γ production by CD8⁺ T cells

Having confirmed that systemic CD8^+^ T cells from OS patients display paralyzed functionality, we next explored the role of gut-associated metabolites (GAMs) in rejuvenating the effector responses. PBMCs isolated from OS patients were treated with representative tryptophan metabolites (HIAA and ILA), bile acids (DCA, 3-Oxo LCA, 7-Keto LCA, and AILCA), and short-chain fatty acids (Acetate, Butyrate, Propionate and Valerate), with cisplatin serving as the reference control of basal level trigger of T cells activation. Among all GAMs screened in the present study, only ILA and AILCA significantly enhanced intracellular IFN-γ yield compared with control (cisplatin-treated) T cells (**Figure 3a,b**). Notably, ILA raised the strongest response, indicating that specific gut-derived metabolites can restore the IFN-γ-producing potential of paralyzed CD8^+^ T cells more effectually than conventional chemotherapeutic agents. These observations identify ILA as a potent immunomodulatory metabolite competent to augment CD8^+^ T-cell activation.

**Figure 3.**
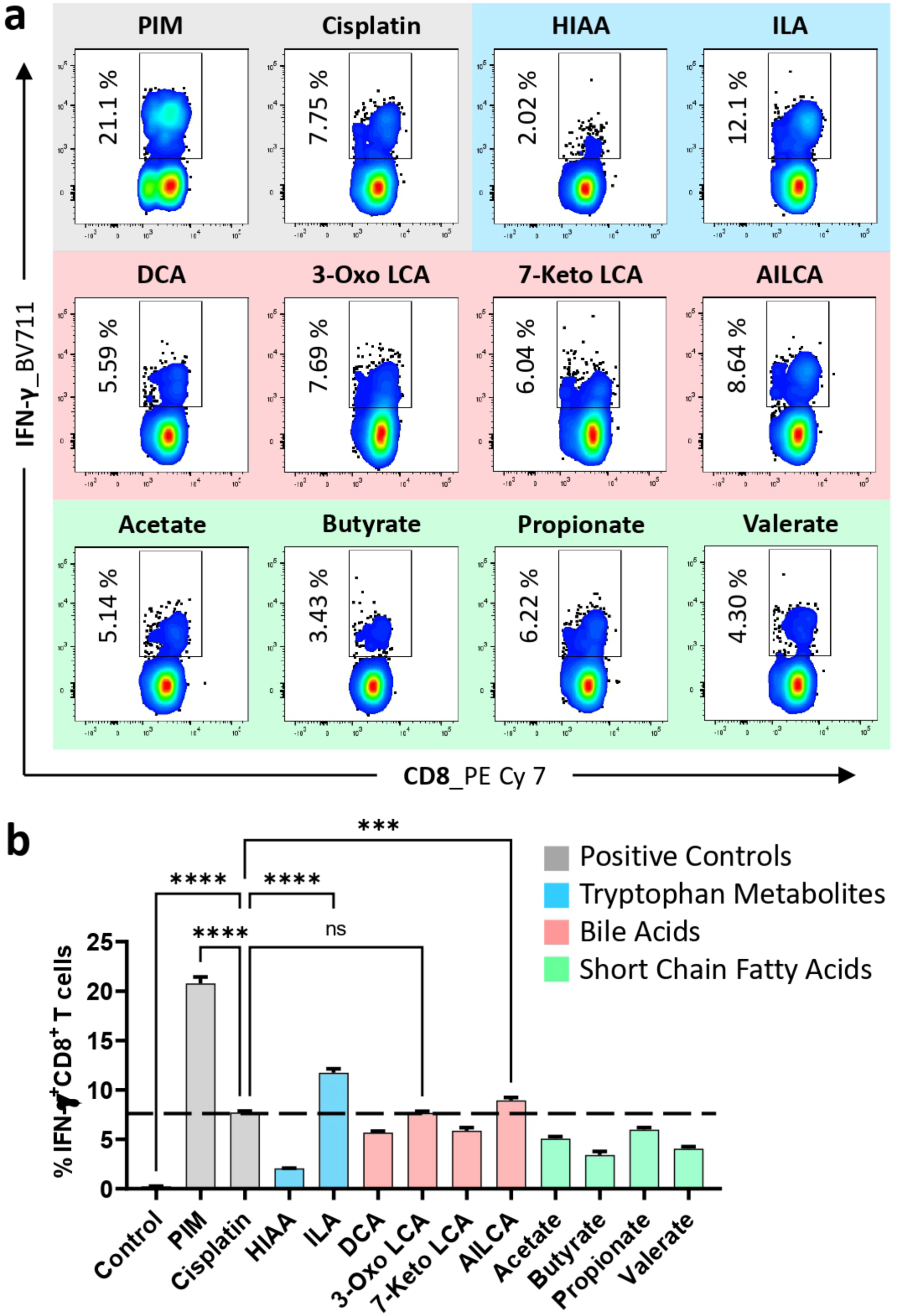
GAMs enhance IFN-γ production by CD8^+^ T cells from osteosarcoma (OS) patients. **(a)** Representative flow cytometry contour plots showing IFN-γ expression in CD8^+^ T cells following 24 h treatment of PBMCs with gut-associated metabolites (GAMs), including tryptophan metabolites [5-hydroxyindoleacetic acid (HIAA) and indole-3-lactic acid (ILA)], bile acids [deoxycholic acid (DCA), 3-oxo lithocholic acid (3-Oxo LCA), 7-keto lithocholic acid (7-Keto LCA), and alloisolithocholic acid (AILCA)], and short-chain fatty acids (acetate, butyrate, propionate, and valerate). PIM (PMA + ionomycin + monensin) served as the positive control for maximal cytokine production, while cisplatin-treated cells were used as a reference for basal T-cell activation. **(b)** Quantification of IFN-γ^+^ CD8^+^ T cells across treatment groups. Among the tested metabolites, only ILA and AILCA significantly increased the frequency of IFN-γ^+^ CD8^+^ T cells above the basal activation threshold. Data are presented as mean ± SEM (n = 3). Statistical analysis was performed using one-way ANOVA followed by Dunnett’s multiple comparisons test. p ≤ 0.05 was considered significant; *p ≤ 0.05, **p ≤ 0.01, ***p ≤ 0.001, and ****p ≤ 0.0001.

### 3.4 GAMs improve TNF-α production by CD8⁺ T cells

Furthermore, to determine whether GAMs largely enhance CD8^+^ T-cell effector function, we next estimated their effects on TNF-α production. Similar to the IFN-γ response, treatment with ILA and Valerate considerably aggravated intracellular TNF-α yield in CD8^+^ T cells at non-cytotoxic concentrations compared with control (cisplatin-treated) CD8^+^ T-cell (**Figure 4a,b**). The potential of these metabolites to augment two independent effector cytokines further mounts their immunostimulatory prospect, with ILA being one of the most compelling metabolites.

**Figure 4.**
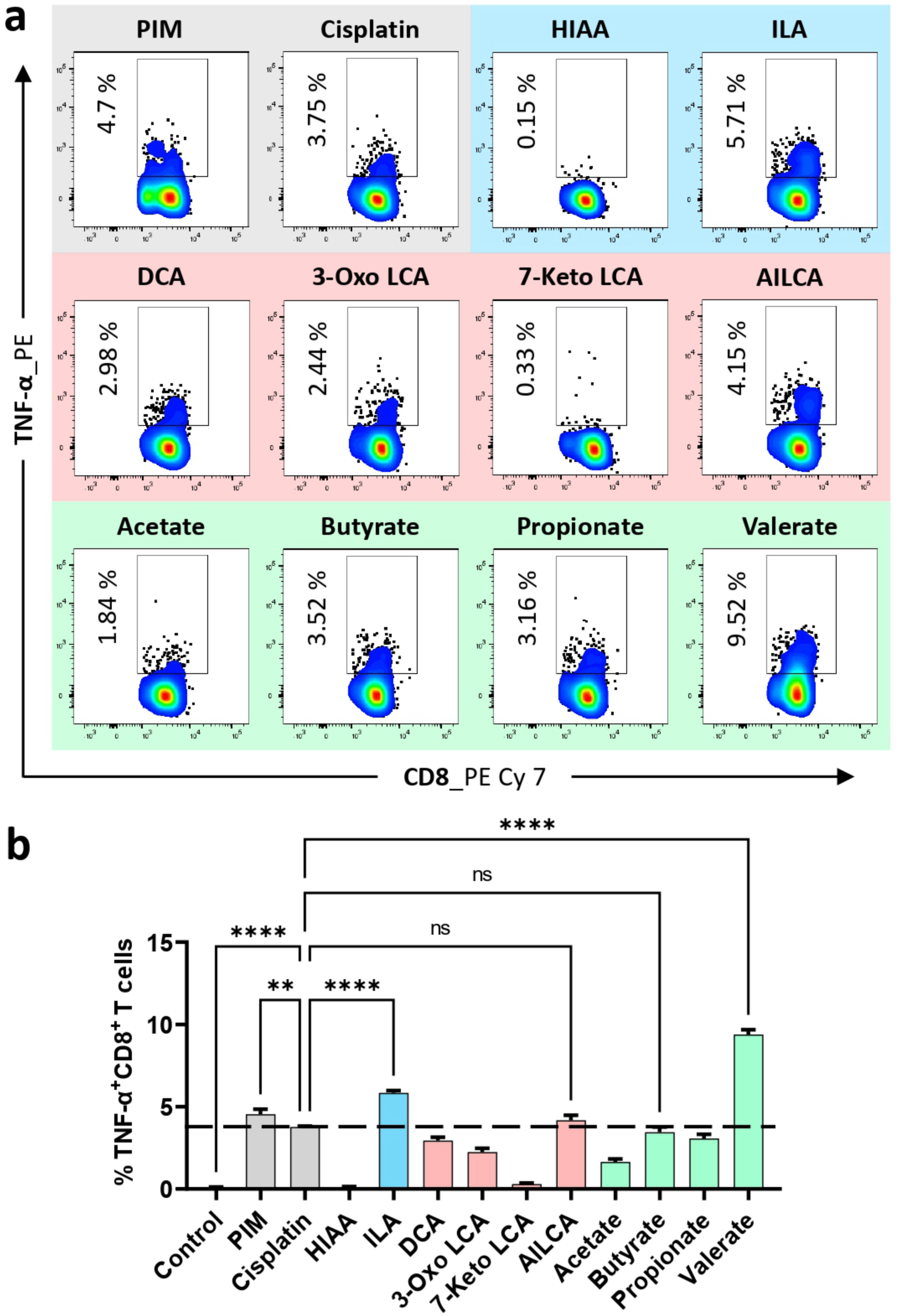
GAMs enhance TNF-α production by CD8^+^ T cells from osteosarcoma (OS) patients. **(a)** Representative flow cytometry contour plots showing TNF-α expression in CD8^+^ T cells following 24 h treatment of PBMCs with gut-associated metabolites (GAMs), including tryptophan metabolites [5-hydroxyindoleacetic acid (HIAA) and indole-3-lactic acid (ILA)], bile acids [deoxycholic acid (DCA), 3-oxo lithocholic acid (3-Oxo LCA), 7-keto lithocholic acid (7-Keto LCA), and alloisolithocholic acid (AILCA)], and short-chain fatty acids (acetate, butyrate, propionate, and valerate). PIM (PMA + ionomycin + monensin) served as the positive control for maximal cytokine production, while cisplatin-treated cells were used as a reference for basal T-cell activation. **(b)** Quantification of TNF-α^+^ CD8^+^ T cells across treatment groups. Among the tested metabolites, only ILA and valerate significantly increased the frequency of TNF-α^+^ CD8^+^ T cells above the basal activation threshold. Data are presented as mean ± SEM (n = 3). Statistical analysis was performed using one-way ANOVA followed by Dunnett’s multiple comparisons test. p ≤ 0.05 was considered significant; *p ≤ 0.05, **p ≤ 0.01, ***p ≤ 0.001, and ****p ≤ 0.0001.

### 3.5 ILA commits the development of polyfunctional CD8⁺ T cells

Polyfunctional CD8^+^ T cells, competent to produce both IFN-γ and TNF-α are widely realized as outstanding mediators of anti-tumor immune responses. We thus next examined whether GAMs treatment could alleviate the proportion of double positive (IFN-γ^+^TNF-α^+^) CD8^+^ T cells. Interestingly, treatment with ILA, AILCA, and Valerate significantly upregulate the percentage of IFN-γ^+^TNF-α^+^ polyfunctional CD8^+^ T cells compared with control (cisplatin-treated) CD8^+^ T-cell, with ILA prompting the most pronounced peak (**Figure 5a,b**). Taken together our current data establishes ILA as the most potent immunomodulator, bolstering polyfunctional CD8^+^ T cells, thereby highlighting its prospective role as a novel therapeutic agent against osteosarcoma. To further validate the polyfunctionality of ILA-primed CD8^+^ T lymphocytes, we next performed gene expression studies for effector cytokines (such as IFN-γ, TNF-α, Perforin-1 and Granzyme-B) using quantitative PCR (qRT-PCR). We observed statistically significant upregulated gene expression of all four effector cytokines in ILA-primed CD8^+^ T cells compared with cisplatin-treated reference control CD8^+^ T cells, reconfirming the polyfunctionality of ILA-primed CD8^+^ T cells **(Supplemental Figure S1b-d)**.

**Figure 5.**
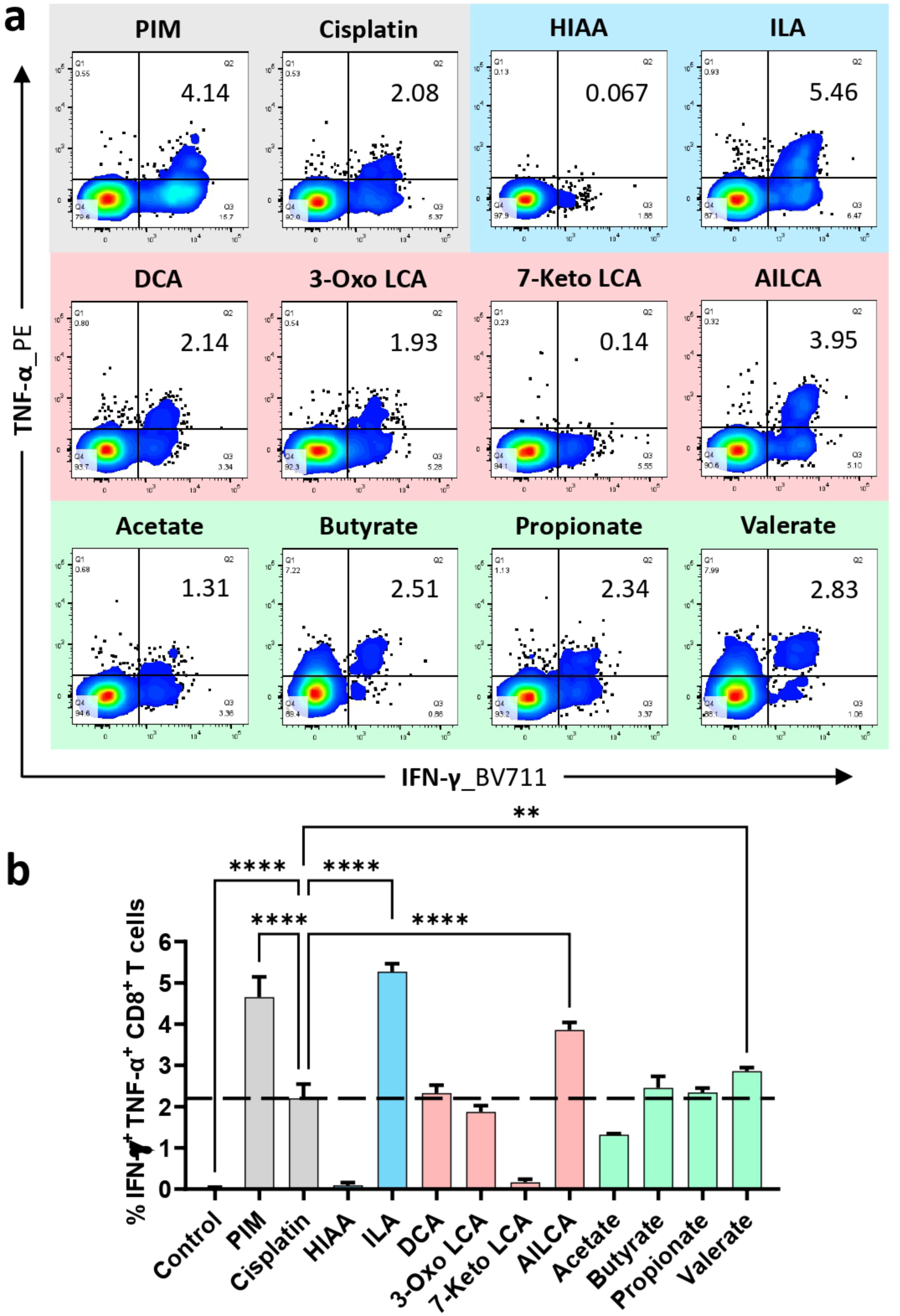
GAMs enhance the polyfunctional anti-tumor activity of CD8^+^ T cells from osteosarcoma (OS) patients. **(a)** Representative flow cytometry contour plots showing IFN-γ^+^TNF-α^+^ CD8^+^ T cells following 24 h treatment of PBMCs with gut-associated metabolites (GAMs), including tryptophan metabolites [5-hydroxyindoleacetic acid (HIAA) and indole-3-lactic acid (ILA)], bile acids [deoxycholic acid (DCA), 3-oxo lithocholic acid (3-Oxo LCA), 7-keto lithocholic acid (7-Keto LCA), and alloisolithocholic acid (AILCA)], and short-chain fatty acids (acetate, butyrate, propionate, and valerate). PIM (PMA + ionomycin + monensin) served as the positive control for maximal cytokine production, while cisplatin-treated cells were used as a reference for basal tumoricidal activation. **(b)** Quantification of IFN-γ^+^TNF-α^+^ CD8^+^ T cells across treatment groups. Among the tested metabolites, ILA, AILCA, and valerate significantly increased the frequency of polyfunctional CD8^+^ T cells above the basal activation threshold, with ILA eliciting the strongest response. Data are presented as mean ± SEM (n = 3). Statistical analysis was performed using one-way ANOVA followed by Dunnett’s multiple comparisons test. p ≤ 0.05 was considered significant; *p ≤ 0.05, **p ≤ 0.01, ***p ≤ 0.001, and ****p ≤ 0.0001.

### 3.6 ILA replenishes the metabolic fitness of CD8⁺ T cells

Given its strong potential to improve the effector functions of CD8^+^ T-cell, we next explored whether ILA could also replenish cellular metabolic fitness. CD8^+^ T cells isolated from OS sufferers were treated with ILA, and cellular uptake of glucose together with mitochondrial bio-mass were examined by flow cytometry. ILA treatment significantly upregulated glucose uptake and expanded mitochondrial bio-mass compared with PIM-treated CD8^+^ T cells, isolated from OS sufferers, to some extent restoring these metabolic parameters toward those noted in healthy controls (**Figure 6a,d**). These findings hint that the immunostimulatory effects of ILA are escorted by ameliorated cellular bioenergetics, furnishing a metabolic foundation for upregulated CD8^+^ T-cell function.

**Figure 6.**
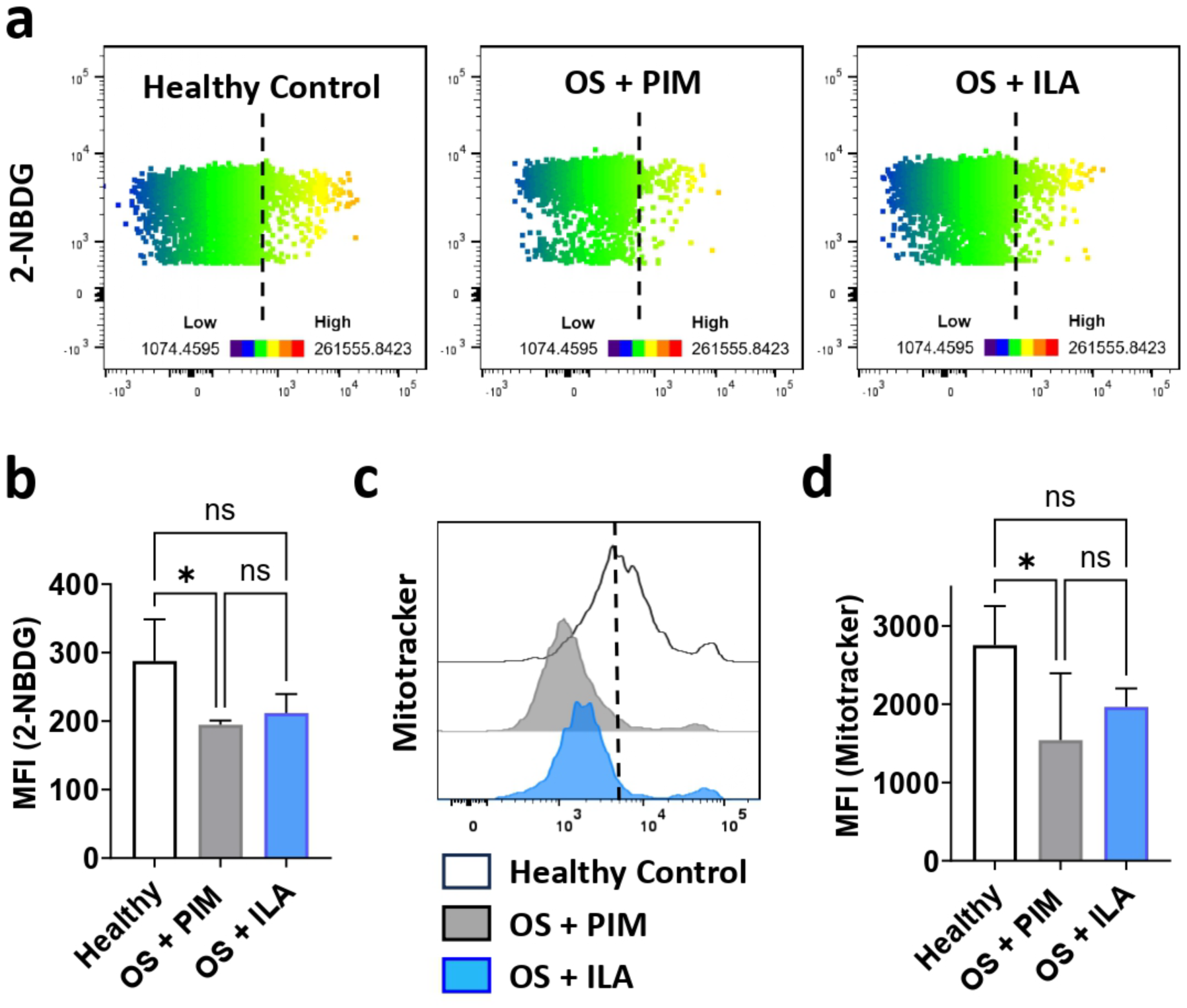
Indole-3-lactic acid (ILA) enhances the metabolic fitness of CD8^+^ T cells from osteosarcoma (OS) patients. Peripheral blood mononuclear cells (PBMCs) from healthy controls and OS patients were cultured under optimal conditions. Healthy control PBMCs and one fraction of OS patient PBMCs were stimulated with PMA (50 ng/mL) and ionomycin (1000 ng/mL) for 5 h, while a separate fraction of OS patient PBMCs was treated with indole-3-lactic acid (ILA) for 24 h. Cells were then stained with 2-NBDG and MitoTracker and analysed by flow cytometry. **(a, b)** Representative flow cytometry plots (heatmap dot plot) and quantification of glucose uptake (2-NBDG) in CD8^+^ T cells. **(c, d)** Representative histograms and quantification of mitochondrial biomass (MitoTracker). ILA treatment significantly improved glucose uptake and mitochondrial biomass in CD8^+^ T cells from OS patients compared with PIM-treated OS patient cells. Data are presented as mean ± SEM (n = 3). Statistical significance was determined using an unpaired Student’s t-test. p ≤ 0.05 was considered significant; p ≤ 0.05 versus the indicated group.

### 3.7 ILA kick-starts CTL-mediated cytotoxicity against osteosarcoma cells

Since ILA invariably displayed the most prominent immunostimulatory activities, we further investigated whether its effects on upregulated cytokine production could be exploited translationally. ILA-primed human CD8^+^ cytotoxic T lymphocytes (CTLs) were co-cultured with U2OS (human osteosarcoma cell-line) at various effector-to-target (E:T) ratios, and apoptosis of tumor cells were quantified by Annexin V/PI counter-staining following flow cytometric inspection (**Figure 7a**). Notably. ILA priming ameliorates CTL-mediated apoptotic induction of target cells (human osteosarcoma cell line, U2OS), compared with PIM-primed CTLs (**Figure 7b**). Quantitative analysis further confirmed significantly upregulated apoptosis of U2OS cells at both 1:1 and 5:1 E:T ratios, with maximal cytotoxicity observed at the 5:1 ratio. Conclusively, our findings establish ILA as a novel immunomodulator, restoring both the cytotoxic potential and cellular metabolic fitness of CTLs, underscoring its candidature for potent therapeutics against osteosarcoma.

**Figure 7.**
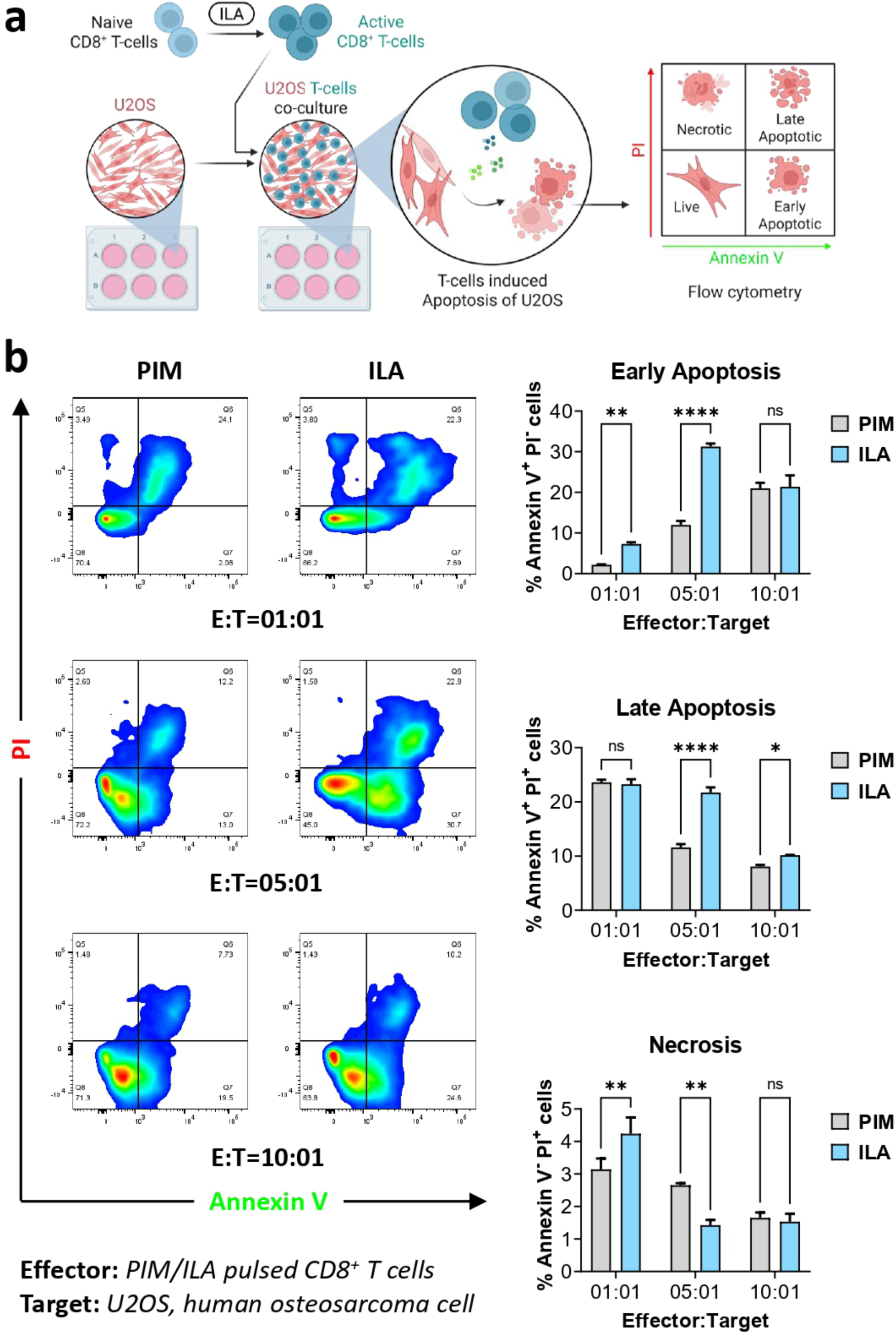
ILA potentiates the cytotoxic function of CTLs against osteosarcoma cells. **(a)** Schematic representation of the experimental workflow for the apoptosis assay. Cytotoxic T lymphocytes (CTLs) were primed with indole-3-lactic acid (ILA) and subsequently co-cultured with U2OS osteosarcoma cells to evaluate CTL-mediated tumor cell killing. Apoptosis of U2OS cells was assessed by flow cytometry. **(b)** Quantitative analysis of U2OS cell apoptosis following co-culture with control or ILA-primed CTLs at effector-to-target (E:T) ratios of 1:1 and 5:1. ILA priming significantly enhanced the tumoricidal activity of CTLs compared to PIM-primed CTLs, resulting in increased apoptosis of U2OS cells at both E:T ratios, with the greatest cytotoxic effect observed at the 5:1 ratio. These findings demonstrate that ILA effectively augments the anti-tumor effector function of CTLs against osteosarcoma cells. Data are presented as mean ± SEM. Statistical significance was determined as indicated; *p ≤ 0.05, **p ≤ 0.01, ***p ≤ 0.001, and ****p ≤ 0.0001.

## Discussion

Osteosarcoma (OS) is depicted by an extremely immunosuppressive “cold” tumor microenvironment, which limits effective anti-tumor immunity and confers a poor prognostic feature^7,12^. Presently, we reveal that circulating CD8^+^ T lymphocytes from osteosarcoma patients exhibit both functional as well as metabolic impairment, emphasizing the close connection between effector immune functionality and cellular metabolism. Emerging studies have confirmed that stable cellular metabolism is an indispensable criterion for sustained activation of CD8^+^ T lymphocytes. Steady state cellular metabolism upregulates cytokine production, cytotoxic potential and tumoricidal potential of T cells^8,9^. However, immunosuppressive, hypoxic and nutrient deficient tumor microenvironment downregulates effector functionality, cytotoxicity and promotes exhaustion of T cells. On similar line, our group identified a novel approach to revert the dysfunctional CD8^+^ T cells to an aggressively cytotoxic, tumoricidal phenotype. We thus activated patient-derived dysfunctional CD8^+^ T cells with an array of GAMs, interestingly Indole-3-lactic acid (ILA) was identified as the most prominent immunomodulatory GAM, inducing the strongest CD8^+^ T cell activation. ILA priming improved polyfunctionality (simultaneous production of multiple effector cytokine, such as IFN-γ, TNF-α, granzyme-B and perforin-1) as well as metabolic fitness, eventually restoring CTLs-mediated killing of osteosarcoma cells. Together, these findings provide a strong foundation for a previously underexplored *“gut-immune-bone tumor”* nexus that could be clinically exploited as a novel therapeutic approach to kick-start anti-tumor immunity against osteosarcoma.

Emerging evidence suggest that gut microbiota and its derivatives (GAMs) influence the prognostic efficacy of immunotherapy against cancer by regulating cellular differentiation, effector function and metabolic status of immune-cells^26–29^. Short-chain fatty acids and indole derivatives have been reported to regulate CD8^+^ cytotoxic T cell function through metabolic and epigenetic reprogramming, thereby modulating anti-tumor immunity^21,30^. Corroborating these observations, our findings suggest that ILA could be utilized as a safe, cost-effective, novel immunomodulatory adjuvant competent to restore dysfunctional CD8^+^ T cell responses against osteosarcoma. Rather than serving as a standalone therapy, ILA could be incorporated in combinational treatment strategies. Thus, ILA or ILA-primed CD8^+^ T cells might further augment the prognostic efficacy of existing therapy, including adjuvant-based chemotherapy, immune checkpoint inhibition or adoptive cell therapy to achieve disease free overall survival.

Since osteosarcoma mainly affects youngsters and adolescents, there is an urgent need for safer, less toxic and tolerable treatment strategies. As a naturally occurring gut microbiota-derived metabolite, especially treatment with ILA may offer a biologically compatible approach to promote anti-tumor immunity alongside existing therapies. However, further preclinical and clinical validations are needed to confirm its safety and therapeutic potential. Nevertheless, the present study has certain limitations. Firstly, due to the rarity of osteosarcoma in youngsters and difficulty in recruiting treatment naive patients, the clinical cohort was small. Secondly, in practice, most patients would have received some form of chemotherapy before their referral. In addition, ethical concerns about blood volume and family reluctance in frail young patients limited sample availability for extensive immune studies in a larger cohort. Furthermore, the present research was limited to *ex-vivo* immunophenotyping and *in-vitro* cytotoxicity assays. Therefore, future validation in larger multicentric cohorts together with mechanistic detailing and preclinical *in-vivo* validation will be integral to establish the effective therapeutic potential of ILA.

In conclusion, our study for the first time uncovered a previously underexplored link between gut-associated metabolites and anti-tumor immunity in osteosarcoma. We discovered the immunomodulatory efficacy of ILA to restore the functional as well as metabolic competence of patient-derived dysfunctional CD8^+^ T lymphocytes against aggressive osteosarcoma cells. These findings provide a strong experimental foundation for the formulation of gut microbiota-derived immunometabolic interventions and support the future evaluation of ILA as a combinatorial adjuvant against osteosarcoma.

## Acknowledgment

LS, AB, AP, AP, TK, SPK, CS and RKS acknowledge the Department of Biotechnology and Central Core Research Facility (CCRF), AIIMS, New Delhi, India, for providing infrastructural facilities. AB and AP thank ICMR for the research fellowship. SPK thanks the DBT for the research fellowship. CS thanks the CCRH-Ayush for the Research Fellowship.

## Credit authorship contribution statement

RKS contributed to the conception and design of the study; LS and AB contributed to the acquisition and analysis of data; AP contributed to data analysis, preparing the figures and the entire manuscript drafting; AF, TK and SPK helped with patient sample handling and preliminary data acquisition; CS helped with RT-PCR; VSK and SA provided human samples. All authors reviewed the manuscript. All authors contributed to the article and approved the submitted version.

**<u>Send correspondence to:</u> Dr. Rupesh K. Srivastava**, Additional Professor, *Translational Immunology, Osteoimmunology & Immunoporosis Lab (TIOIL) An ICMR Collaborating Centre of Excellence on Bone Health* Department of Biotechnology, All India Institute of Medical Sciences (AIIMS), New Delhi-110029, India Cell: +91-9179567399

---

**Supplemental Figure S1.**
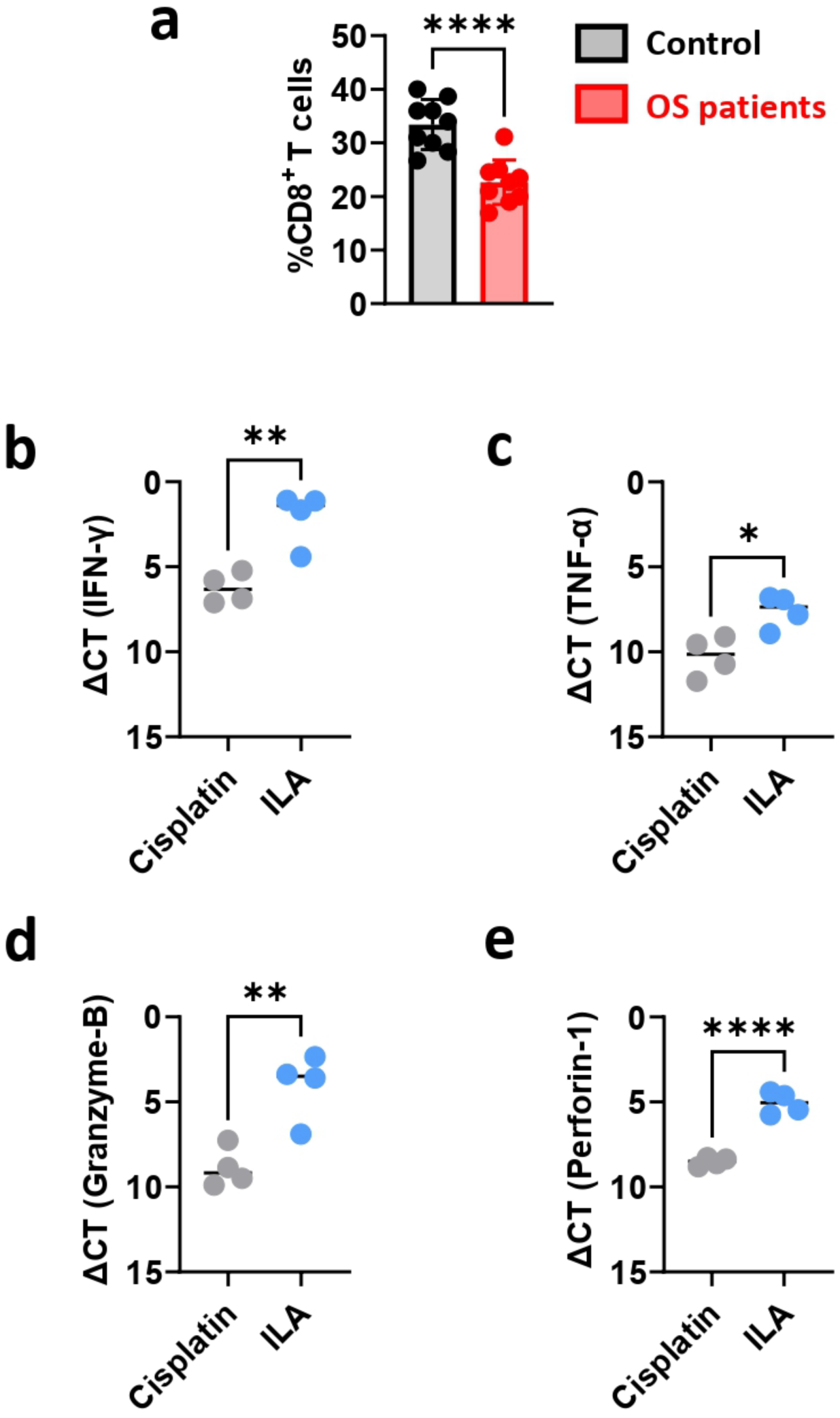
Reduced CD8^+^ T cell frequency in OS patients and ILA-mediated enhancement of effector gene expression in OS patient-derived CD8^+^ T cells. **(a)** Frequency of circulating CD8^+^ T cells in peripheral blood of healthy controls and osteosarcoma (OS) patients, determined by flow cytometry. **(b-e)** Quantitative RT-PCR analysis of IFN-γ, TNF-α, Granzyme-B, and Perforin-1 expression in CD8^+^ T cells isolated from OS patients following stimulation with ILA or cisplatin (reference control). Data are presented as mean ± SEM. Statistical significance was determined using an unpaired Student’s t-test (*p ≤ 0.05, **p ≤ 0.01, ***p ≤ 0.001, and ****p ≤ 0.0001).

